# On the relationship between spatial environmental variability, dispersion and biodiversity

**DOI:** 10.1101/2024.08.15.608079

**Authors:** Giulia Bernardini, Gareth G. Roberts, Mark Sutton

## Abstract

**Aim:** We establish quantitative relationships between species richness and the rate of spatial change in controlled, digital, environments. We use a simplified, first-principles, stochastic, evolutionary model in which artificial organisms can evolve and disperse. We develop an understanding of how environmental variability in space influences species richness and how it is affected by organisms’ ability to disperse.

**Time period:** At each time step of the experiment, each organism can reproduce sexually, disperse and die. Each experiment is run for 100,000 time steps. The life span of each organism is 15 time steps.

**Location:** The model uses an artificial, digital, landscape consisting of a uniform (*x, y*) grid of cells, with a single environmental variable that changes sinusoidally in the *x* direction.

**Major taxa studied:** Organisms are defined by a 64-bit genome and reproduce sexually. These digital organisms are designed to mimic the basic principles of biological evolution.

**Methods:** Each experiment starts with a single organism, which can mutate and reproduce sexually, producing offspring that can do all of the above and disperse in the environment. Monte Carlo experimentation is used to generate statistical insight into species richness by producing thousands of replicate simulations.

**Results:** A strong correlation is observed between mean species richness and the rate of spatial change in the environmental variable. This relationship holds true for a wide range of dispersal abilities, but diminishes when dispersal ability is very low.

**Main Conclusions:** We predict that the rate of change of environmental variables (e.g. the derivative of elevation) is a driver of real-world biodiversity where dispersal is sufficient. Our results suggest that the ability of organisms to disperse plays an important role in determining how biodiversity responds to environmental gradients.

## 1 Introduction

Understanding the drivers of biodiversity remains a central challenge in ecological research (e.g. Mitchell et al., 2024). It is generally accepted that distributions of species in space and time depend upon abiotic (external) and biotic drivers (e.g. Franklin, 2010; Guisan et al., 2017). Establishing correlations (or coherence) between environments and species richness (species within a particular area) has been greatly facilitated by the availability of modern expert range maps, atlases and palaeobiological datasets (e.g. GBIF, PBDB; Jenkins et al., 2013, 2015; O’Malley et al., 2023). This work has emphasised the importance of abiotic drivers of species richness including spatial variability of climate, elevation and temperature (e.g. Antonelli et al., 2018).

It has also helped to establish the importance of scale in determining strength of coherence between abiotic processes and biodiversity (e.g. Belmaker and Jetz, 2011; O’Malley et al., 2023). The proliferation of Machine Learning methodologies in macroecology has focused attention on identifying statistical relationships between species richness, mapped using atlas data for instance, and environmental variables (e.g. Phillips et al., 2006; Liu et al., 2024). In contrast with those studies, we explore whether observed correlation between environment and species richness can be explained by theory that incorporates evolution’s natural stochasticity (e.g. Yaeger, 1994; Garwood et al., 2019; Dolson and Ofria, 2021; Furness et al., 2021).

This study aims to develop an understanding of how spatial rates of environmental change relate to species richness (Figure 1). We further examine how this relationship varies when the capability of organisms to disperse is varied. We utilise the Rapid Evolutionary Simulator (REvoSim), which implements an efficient organism-level ‘first principles’ eco-evolutionary system incorporating spatial structure. In this system species emerge ‘naturally’, and hence distributions of species are generated in response to deterministic environmental forcing and stochastic evolution (Figure 2a-e; Garwood et al., 2019; Furness et al., 2021). A benefit to adopting such an approach is that generally accepted mechanisms for driving speciation can be isolated and their impact assessed. We seek to establish whether rules-of-thumb, relating environmental spatial variability to distributions of species richness, can be determined whilst embracing the inherently stochastic nature of evolution.

**Figure 1:**
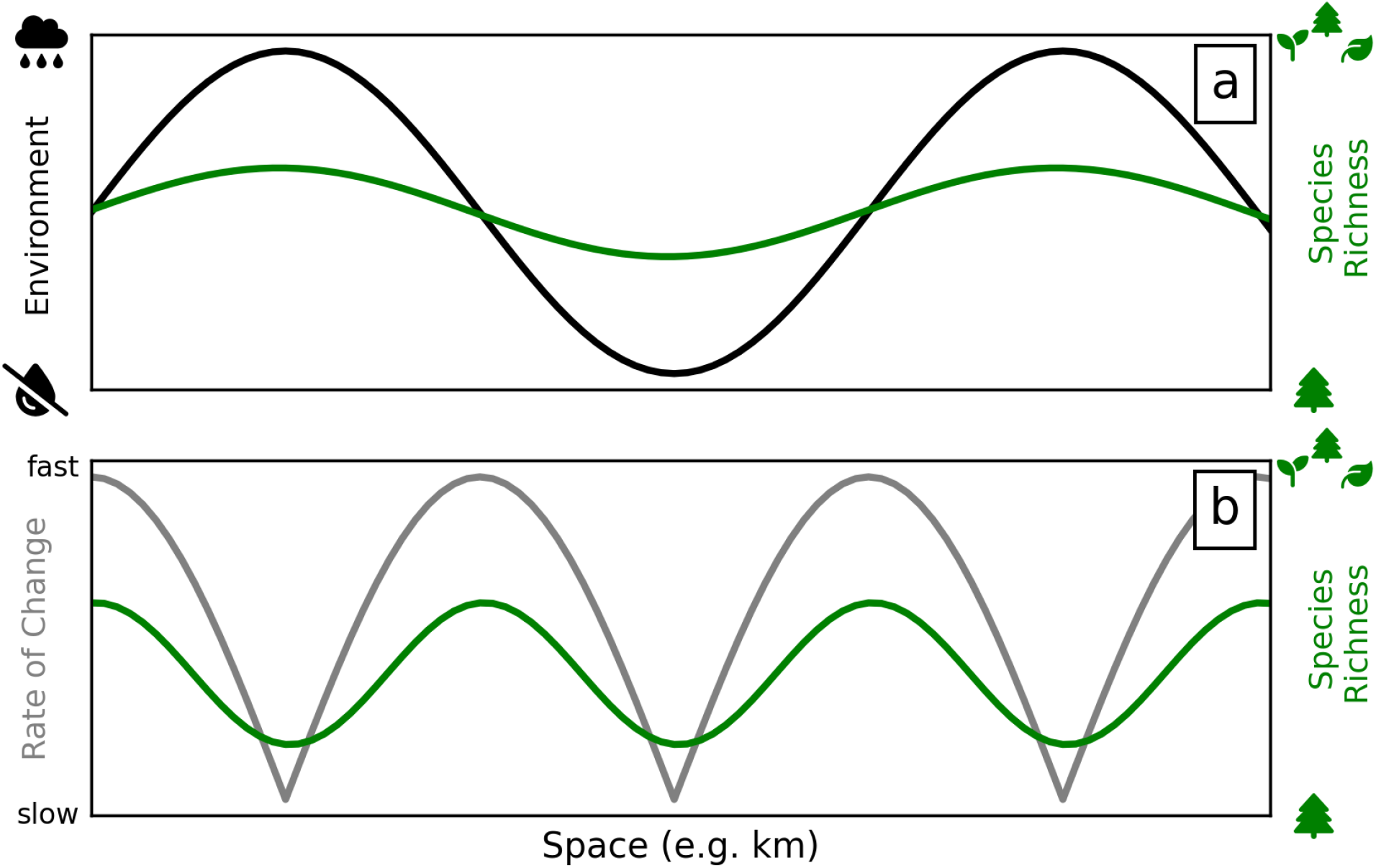
Simplified postulated relationships between environment and species richness tested in this study. (a) Species richness (green) has a simple and direct relationship with environment (black). More formally, species richness is in-phase with the environment. (b) Species richness (green) has simple and direct relationship to environmental rate of change (gray). Strictly, we consider the absolute value of the derivative of the environmental variable (cf. black and gray curves).

**Figure 2:**
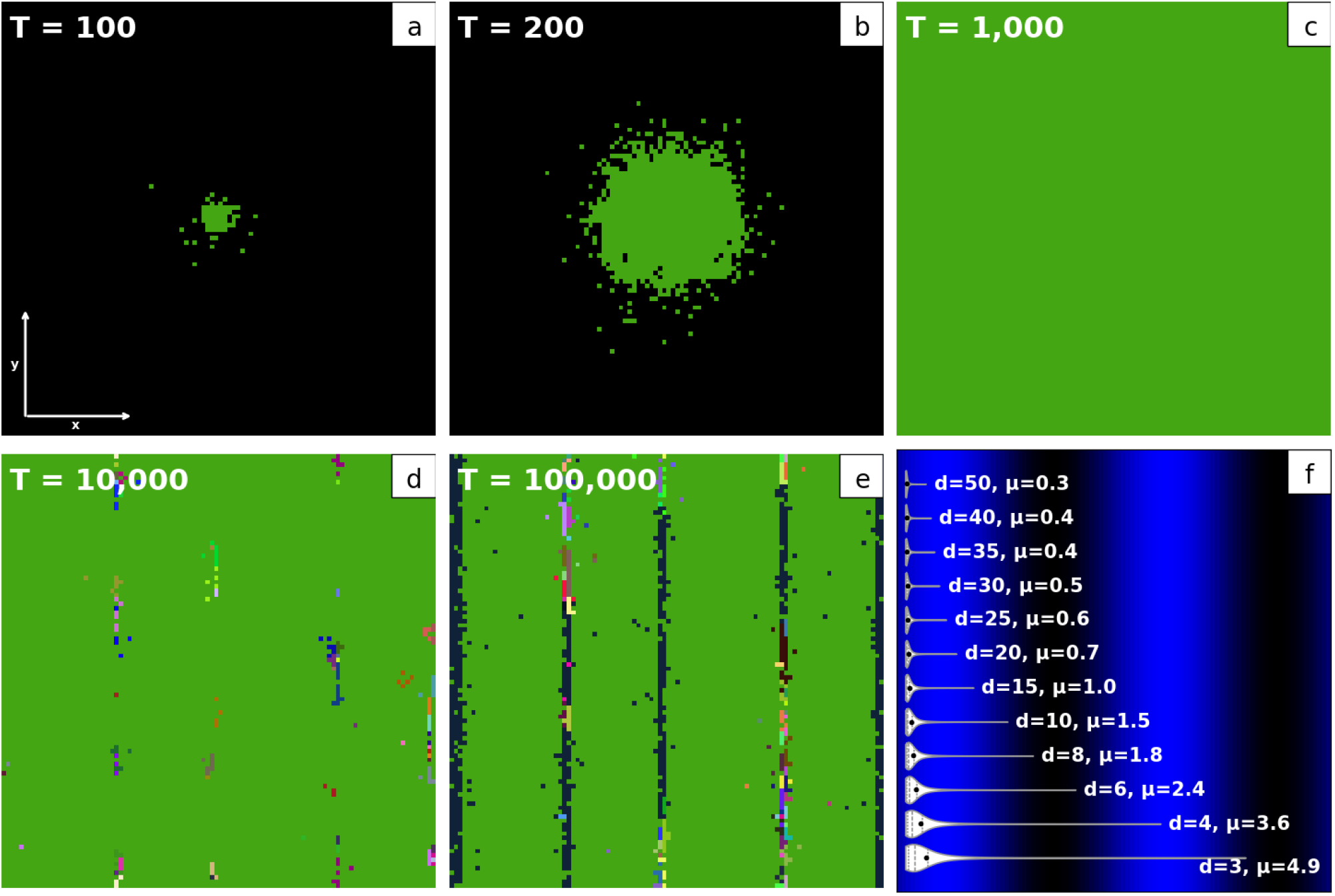
An example of species dispersion in a single simulation and dispersion parameters included in this study. (a-e) Species richness in a single simulation after 100, 200, 1000, 10,000 and 100,000 steps. This simulation has spatial dimensions of 100*×*100 pixels (white vectors in panel a; see body text for extended description). Colours correspond to the most abundant species in a pixel at any given time; colours for species are chosen arbitrarily. Black indicates no organisms present. (a-b) The first organism (green), inserted into the centre of the domain, reproduces and disperses to other pixels. (c) Colonisation of the whole domain. (d-e) Different colours show that speciation has occurred and new species are now most abundant in certain pixels. (f) Map showing the simple sinusoidal environment and dispersion parameters, *d*, used to control dispersion of offspring in this study. White violin plots show probabilities of dispersion for each dispersion parameter at the scale of the environment; black dots = mean dispersion, *µ. d* = 15 for simulation shown in panels (a–e). See body text for details.

## 2 Methods

### 2.1 Modelling strategy

REvoSim simulates evolution of artificial organisms within environments that can vary over space and time (Garwood et al., 2019). The organisms can reproduce and mutate. Each individual is defined by a 64-bit binary genome, a sequence of zeros and ones (genes). Genomes within populations evolve through stochastic mutation; genes are also subject to adaptation through selection for fitness to the environment. Species are automatically indentified by REvoSim using the Biological Species Concept, i.e. as reproductively-isolated population groups (Garwood et al., 2019). For simplicity, we use species richness as a measure of biodiversity. This metric has been criticised as it does not account for relative abundance, genetic variation, or functional traits (see e.g., Gotelli and Colwell, 2001; Hillebrand et al., 2018); it is nonetheless widely used as it is relatively easy to determine in real-world datasets. In order to determine whether choice of metric influences our results, we compare results generated using species richness with those obtained from genetic diversity measures (Figure S2).

We explore simple models in which environment, *e*, is determined by integer values, 0 ≤ *e* ≤ 255, in a uniform grid of *X × Y* cells or pixels (px), where *X* = *Y* = 100. REvoSim maps environmental parameter values to colours in a raster image for visualisation purposes. We use an environmental map that varies sinusoidally in the *x* direction and is constant in the *y* direction. This setup is analogous to real-world scenarios where organisms must adapt to a single environmental variable that changes along a single spatial dimension. We set the spatial grid to be toroidal, i.e. to ‘wrap around’ in both *x* and *y* directions, allowing the elimination of edge-effects (Porensky and Young, 2013). The sinusoidal wavelength chosen (*λ* = 50 px) provides this ‘wrap around’ in the *x* direction without discontinuities (Figure 2f).

In a series of systematic experiments, we varied the capability of organisms to disperse. In all experiments, the environment and dispersion ability were first defined; other settings were left at REvoSim defaults. At the start of each experiment, a population of genetically identical organisms were inserted into the cell at the centre of the environment, where they started reproduction, mutation, and dispersion. The experiment were then iterated for *T* = 100, 000 steps. Species richness broadly increased in each experiment until ∼ 50, 000 steps, after which it tended to stabilise (Figures 2a-e & 3). *N* = 3000 experiment replicates were produced for each scenario (i.e. for each dispersal setting). This Monte Carlo experimentation allows us to develop an understanding of expected species richness from models incorporating evolutionary stochasticity, which can generate very different trajectories across replicates (Figure 3).

**Figure 3:**
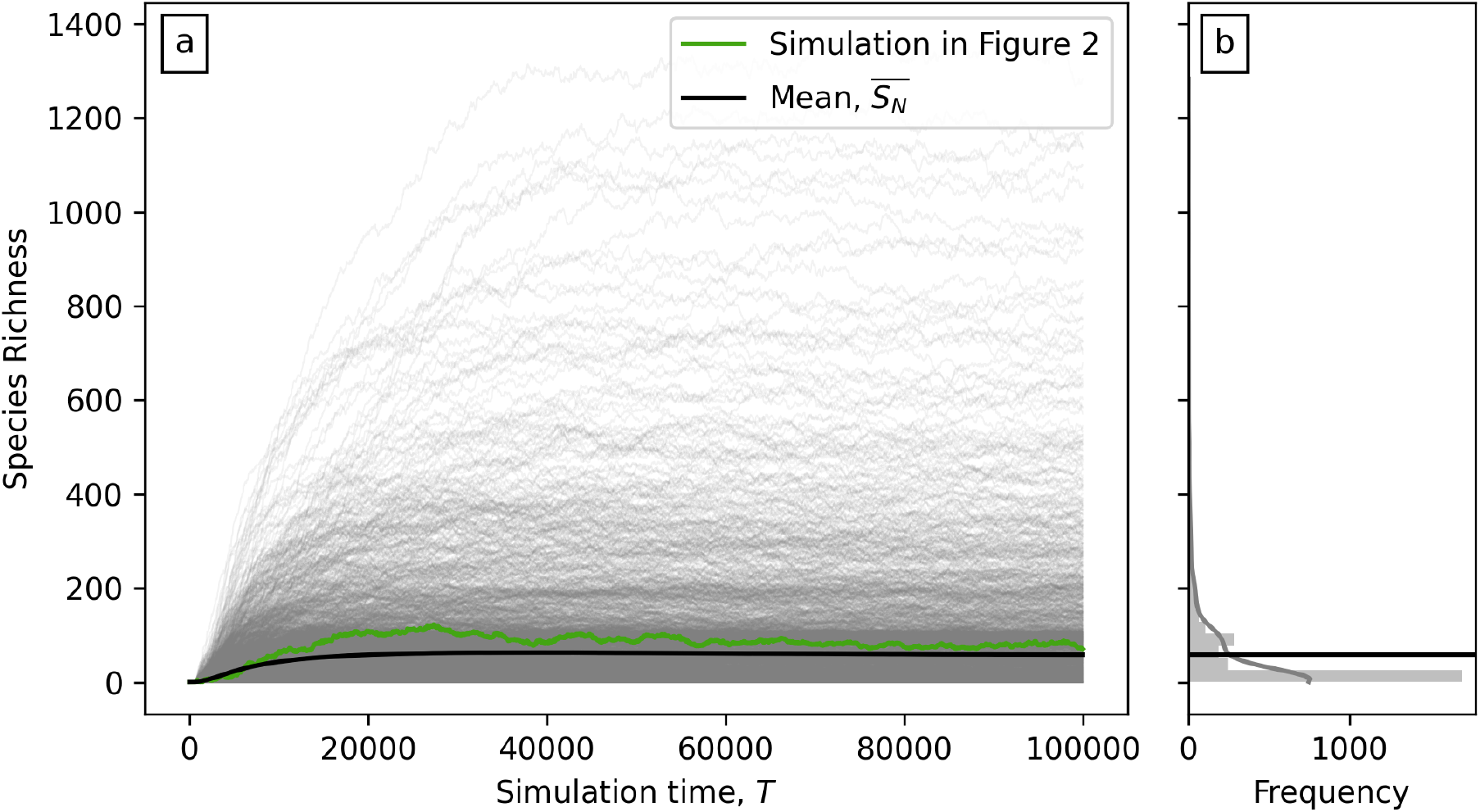
Variability of species richness in time from Monte Carlo experimentation. (a) Gray curves = species richness as a function of time in 3000 simulations. Environmental variable (*λ* = 50 px) and dispersion parameter (*d* = 15; see Figure 1f) were held constant. Gene mutation, reproduction and dispersion are stochastic (see body text for extended description). Green line = trajectory of simulation depicted in Figure 1a–e. Black line = mean species richness, 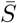, from the 3000 simulations. (b) Gray histogram and black line = distribution of species richness and 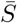 at *T* = 100, 000, respectively.

### 2.2 Evolutionary stochasticity

The replicates incorporated four sources of stochasticity. First, the genome of the initial population is randomly selected from the set of organisms able to survive in the central cell. Secondly, the fitness of each genome to a particular environment is controlled by a ‘fitness landscape’ (McGhee, 2006; McCandlish, 2011), which is randomly generated at the beginning of each replicate (see Garwood et al., 2019). Thirdly, genetic bits can mutate stochastically during reproduction. Dispersal is the fourth stochastic element. Organisms do not move during their lifespan, but offspring of reproductive events ‘disperse’, i.e. attempt to establishing themselves either in the same pixel as their parents or in a nearby pixel. The dispersal distance is defined as

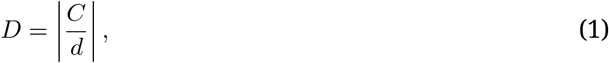

where the dispersion parameter, *d*, is set at the beginning of each experiment and is constant for all organisms. The parameter 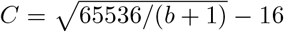 controls the stochasticity of dispersion via the value of *b*, which is chosen randomly from a uniform distribution 0 ≤ *b* ≤ 255 (thus 0 ≤ *C* ≤ 240). In practice, displacement is determined by defining translation along the *x* and *y* axes

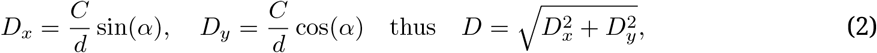

with the azimuth of displacement, 0 *< α* ≤ 360, chosen randomly from a uniform distribution (Garwood et al., 2019).

The dispersion of an organism is inversely proportional to *d*, with minimum and maximum values of zero px and *D*_*max*_ = 240*/d* px, respectively. The distribution of dispersal distances is weighted towards low values. The values of *d* used in this study, along with associated dispersion distributions and mean dispersion, *µ*, are given in Figure 2f. Figure 2a-c shows an example of species spreading across the model domain (via reproduction and dispersal), during 1000 time steps. In this example *d* = 15, and the probability that offspring will disperse to within *D* = 2 px of the pixel inhabited by their progenitors is 88%.

Figure 3a shows species richness trajectories for each of the 3000 replicates produced with *d* = 15. Mean species richness for all replicates (black curve) and the experiment shown in Figure 2a-e (green curve) are indicated. The histogram in Figure 3b shows the distribution of species richness at for all replicates (*N* = 2871) that had extant species at *T* = 100, 000.

### 2.3 Metrics of species richness and genetic diversity

Two types of logs were generated for each individual experiment. The first type records species richness every 50 time steps (referred to as ‘species counts’ by REvoSim). These were configured as ‘v2.0.0 CSV’ logs in REvoSim settings. The second type records the genomes, species identifications, and spatial locations for all individuals present at the final step (*T* = 100, 000). The first type were used to extract species richness across the grid. The latter were used to calculate mean species richness at each pixel and to calculate genetic diversity across the grid.

Species richness for each pixel in each replicate was calculated by counting the unique number of species present in that pixel. These results are used to calculate mean species richness at each *x* position on the grid, for each value of *d* (see Figure 4f). Species richness across the domain was also calculated for each replicate by counting the unique number of species present in the whole 100 *×* 100 pixels grid. These values were then compared with the genetic diversity of the corresponding replicate.

**Figure 4:**
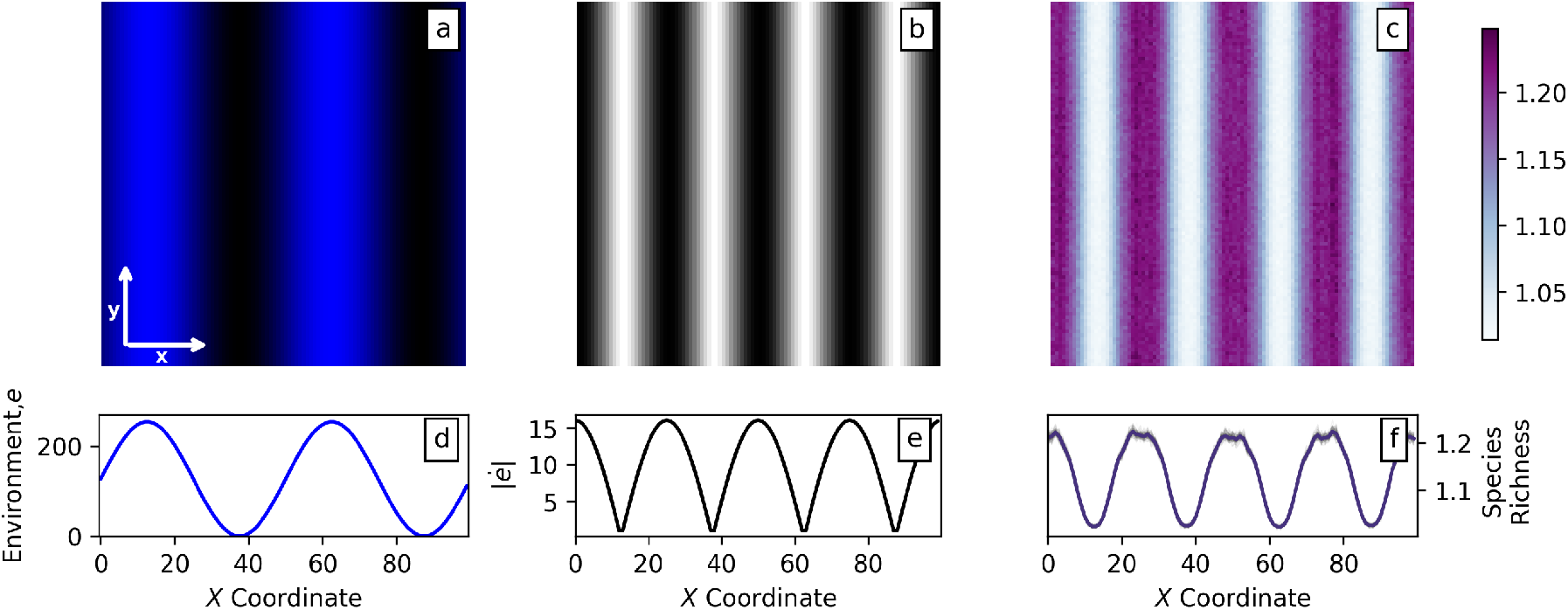
Distribution of species richness at steady-state from Monte Carlo experimentation. (a) Environment; blue/black = high/low values; *λ* = 50 px. (b) Absolute values of the derivative of the environment; white/black = low/high. (c) Mean species richness for each pixel at *T* = 100, 000 steps from the 3,000 simulations (*d* = 15; see Figure 2f). (d) and (e) transects across the centre of the the environment (panel a; at *y* = 50) and the absolute values of its derivative (b), respectively. (f) Transects across mean species richness shown in panel (c); gray curves = transects at *y* = 1, …, 100; purple curve = mean species richness for all 100 transects.

For clarity, we used the following terminology to describe species richness. *S*(*x, y*) refers to the species richness of cell (*x, y*) for a particular value of *d*, in a particular replicate *n*. See Figure S1 for a schematic of the model setup and associated statistics. All of the following statistics were calculated using only the *N* replicates that had extant species at *T* = 100, 000. In practice more than 92% of all simulations are included in these results.

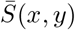 is the mean species richness of cell (*x, y*) for a particular value of *d. S*_*n*_ is the number of unique species across the entire grid for an individual replicate, *n*. 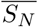 is the mean number of species across the entire grid for a particular value of *d*, over all replicates, 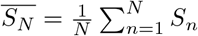.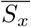 is the mean species richness as a function of *x*, for a particular value of *d* incorporating all replicates. This value is calculated as the mean of 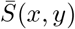 for a given *x*. For example, at *x* = 50, 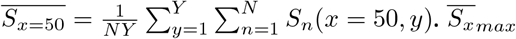 is the maximum value of 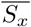 for a specific value of *y* and *d*. Additionally, we calculated the normalised value of mean species richness as a function of *x* as 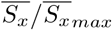. This normalised value is used to compare 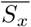 across the different values of *d*.

Mean genetic diversity in a single replicate, 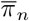, was calculated by comparing genomes from *M* pairs of individuals,

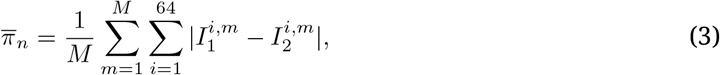

where *I*_1_ and *I*_2_ are the individuals whose genomes were compared, and *i* = 1, 2, …, 64 is the index of genes for each genome for each individual. Since genes are either expressed as a one or a zero when all individuals are genetically identical 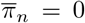. For theoretical scenarios in which all individuals have completely different genomes 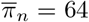. It is possible to calculate the true mean genetic diversity for each replicate. However, given that the number of individuals present during each replicate can exceed *I* = 100, 000, the number of all possible unique pairs to be compared can easily exceed 10 billion (i.e. *I*(*I* − 1)*/*2), which is computationally impractical. Instead, we estimated 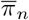 from a sample of the total number of pairings, by comparing *M* = *I/*2 randomly chosen unique pairs. Thus, we typically we compared *>* 90, 000 pairs of individuals for each replicate. Figure S2 plots mean genetic diversity for each replicate, 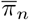 compared with its respective species richness, *S*_*n*_.

## 3 Results

### 3.1 Mean species richness varies with dispersal ability

Figure S2 shows that genetic diversity and species richness covary positively when *d <* 40 (*D*_*max*_ ≥ 6, *µ* ≥ 0.4). When *d* ≥ 40, the absence of visually detectable covariance is a reflection simply of the narrow range of biodiversity variation (in both metrics), resulting in domination by stochastic noise. As species richness is a good proxy for genetic diversity (and is simpler to calculate, both in our replicates and in real-world data), we use it as our biodiversity metric in all analyses.

Species richness for the entire model domain varied between 1 (only one species present) to 10,000 (all pixels occupied by a different species) over the full range of experiments. The highest mean species richness, 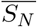, at the final step of the model occurred at *d* = 50 (*D*_*max*_ = 4.8 px, *µ* = 0.3 px; see Figure S4. Figures 3b and S3 show that frequency distributions of species richness tend to be skewed towards low values for high dispersal (*d* ≤ 20, *D*_*max*_ ≥ 12 px, *µ* ≥ 0.7 px), and to be Gaussian or slightly skewed towards higher values for low dispersal (*d* ≥ 25, *D*_*max*_ ≤ 10 px, *µ* ≤ 0.6 px).

### 3.2 Relationship between diversity and environmental parameter

Figure 4 illustrates the spatial distribution of the environmental parameter, the absolute value of its derivative, |*ė*| (= |d*e/*d*x*|), and mean species richness, 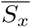, for a representative value of *d* = 15. Visual inspection of these results shows that species richness is highly correlated with the rate of spatial change of the environmental parameter, |*ė*|, rather than with *e* itself. Normalised results for other values of *d* are shown in Figure 5. For clarity, |*ė*| was normalised by dividing by its maximum value such that 0 ≤ |*ė*|*/*|*ė*|_*max*_ ≤ 1. The highest value of value of 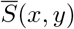 was 2.78, recorded for *d* = 35 (*D*_*max*_ = 8 px, *µ* = 0.5 px).

**Figure 5:**
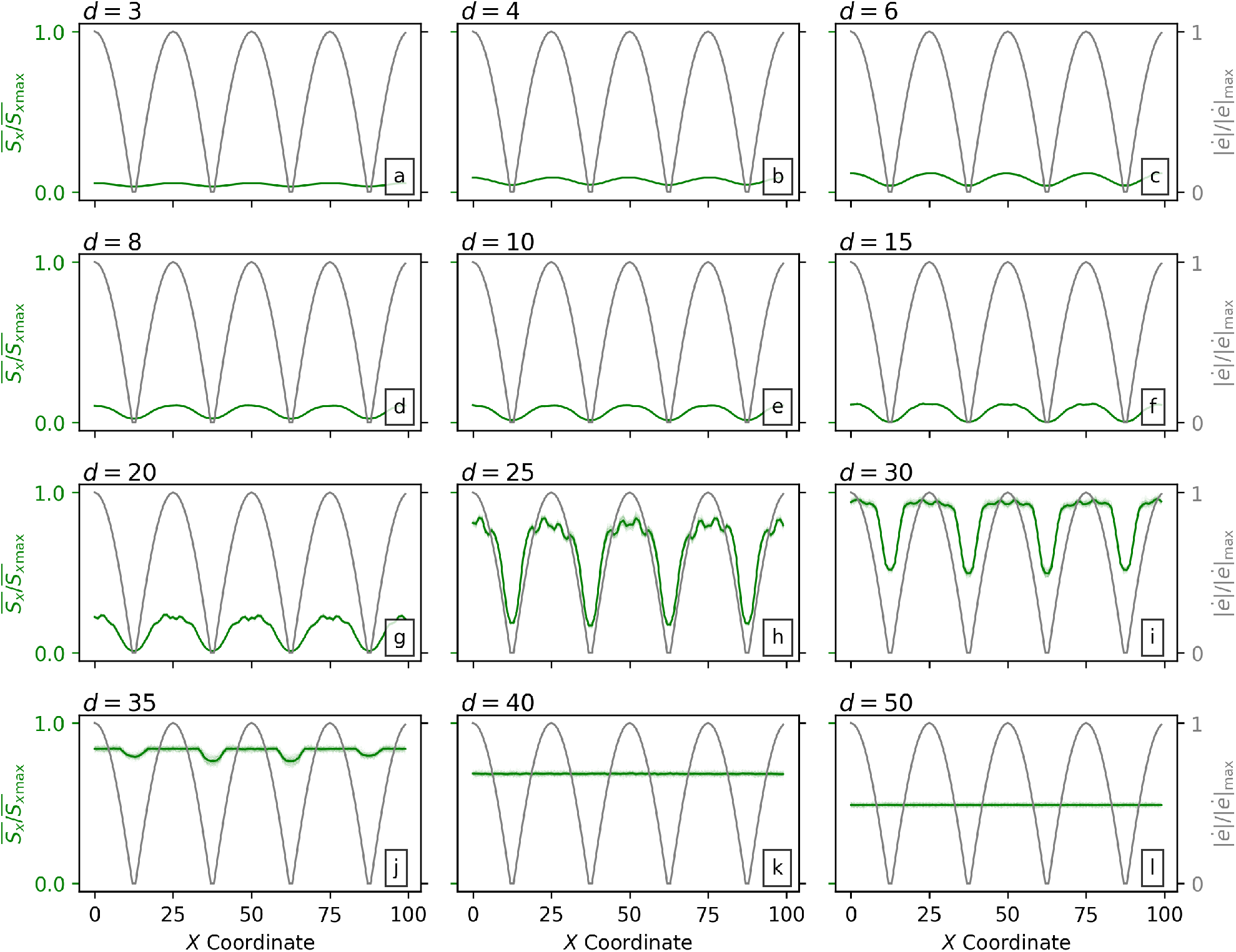
Relationships between environment and species richness as a function of dispersion. (a–l) All simulations used the same environment (see Figure 4; *λ* = 50 px). Dispersion ability was varied by systematically changing the value of *d*, see panel annotations and Figure 2f. Gray curves = absolute value of the derivative of the environmental variable, |*ė*|, normalised by the maximum value, |*ė*|_max_, as a function of *x* coordinate (see Figure 4e). Dark green curves = transects of mean species richness, 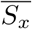, (along the *x* coordinate; see Figure 4f) for 3,000 simulations with given value of *d*, normalised by the maximum mean value of species richness from all 36,000 simulations, 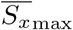. Light green curves = transects at *y* = 1, …, 100; note that in many cases variation is less than the thickness of dark green curve, and hence light green curves are not visible.

For each value of *d*, Pearson’s product-moment correlation coefficients, *r*, were calculated between 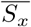 and |*ė*|. The results are shown in Figure 6. High correlations (*r >* 0.8) were observed for *d <* 35 (*D*_*max*_ *>* 7 px, *µ >* 0.4 px; cf. Figure 5).

**Figure 6:**
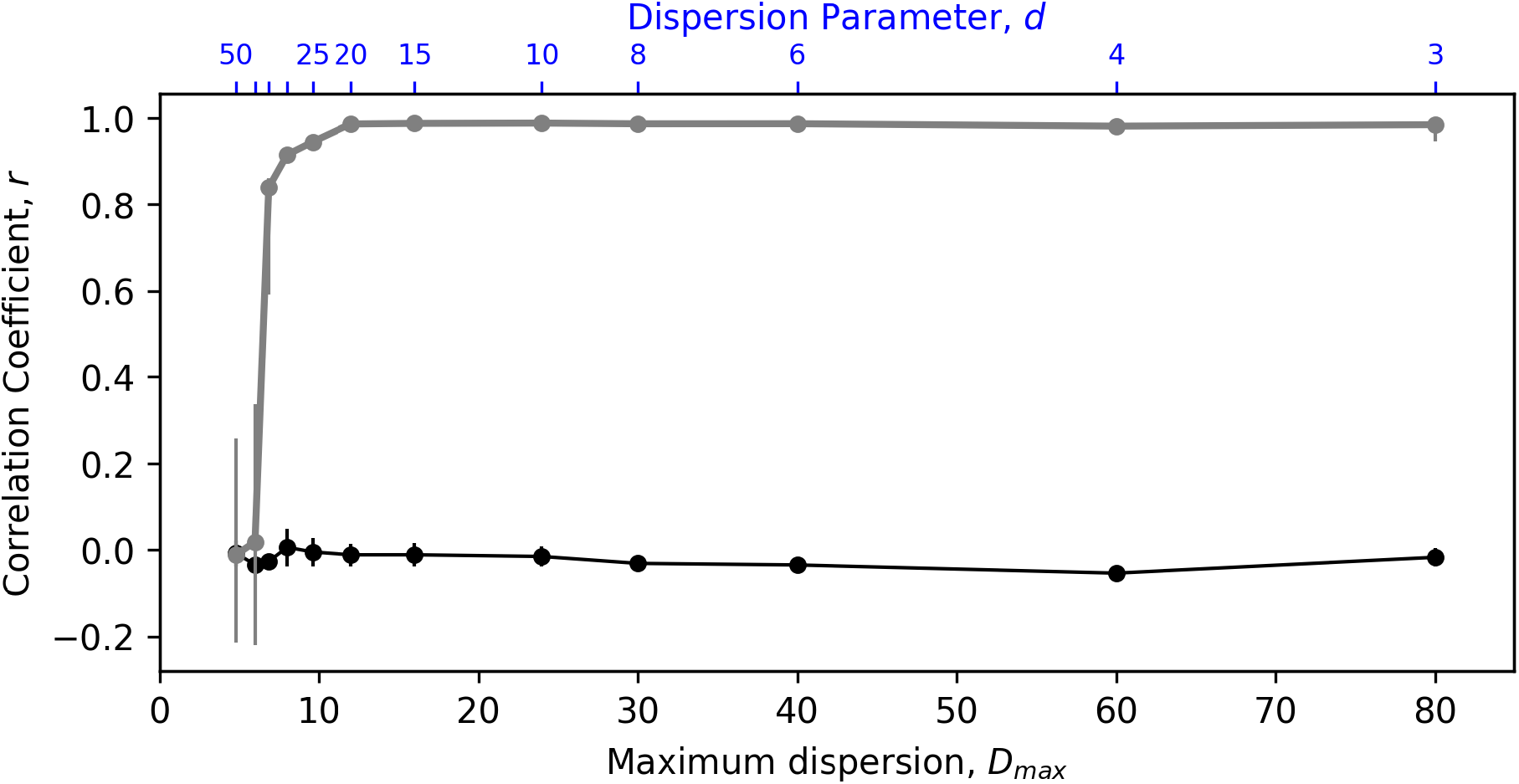
Correlation between species richness, derivative of the environment, and environment as a function of dispersion. Grey circles = Pearson’s correlation coefficients calculated using mean values of species richness, 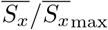, and spatial rate of environmental change |*ė*|*/*|*ė*|_max_, see respective dark green and grey curves in Figure 5. Correlation coefficients are shown for simulations with different dispersion parameter, *d*, values; note that maximum dispersion, *D*_*max*_ (with units px), is annotated (see Figure 2f for details). Error bars = range of coefficients calculated for all, *y* = 1, …, 100, transects in each systematic test of *d* shown in Figure 5. Black circles/error bars = correlation coefficients and ranges for environment and species richness.

## 4 Discussion

### 4.1 Spatial rate of change, species richness and the role of dispersion

Species richness in our theoretical ecosystem, which incorporates stochastic mutation, dispersion and fitness, correlates positively with the absolute value of the derivative of the environment, |*ė*|. In other words, the pace of spatial variation of an environment appears to be a useful predictor of the number of species within a given area when the probability of dispersion is sufficiently high. In this section, we discuss this correlation and its relationship to the adaptation of species to the rate of change of the environment. We first discuss the results in terms of gene flow (*sensu Slatkin, 1987*). Secondly, we discuss the impact of ability to disperse on correlations between species richness and environmental variability. Thirdly, the impact of only permitting the environment to vary in one direction is discussed. Finally, we explore a simple rule of thumb for identifying when environmental conditions and species richness are likely to be highly correlated. In the final section we briefly explore the relevance of this work for understanding relationships between species richness and the environment in the real world.

In Figure 7, the regions (both with spatial range Δ*x*) labelled *i* and *ii* represents areas with low and high environmental variability (Δ*e*), respectively. In other words, the environmental conditions in cells are more similar to their near neighbours in region *i* than in region *ii*. As a consequence, offspring of parents living in cells in region *i* are more likely be adapted to nearby cells, and thus be able to successfully settle in them. Consequently, gene flow, which requires successful settling in non-parental cells, is greater in region *i* than in *ii*. Gene flow between cells tends to homogenise the genomes of populations; adaptation to their local environments tends to differentiate them, eventually into different species (Coyne et al., 2004). The balance between these ‘pressures’ appears to determine the prevalence of speciation, which will be favoured in scenarios where gene-flow is inhibited or adaptive pressures (e.g. arising from environmental variation) are high. As the environmental difference between nearby cells in region *ii* (i.e. high |*ė*|) is greater than that in region *i* (i.e. low |*ė*|), adaptive differentiation pressures between cells is likely to be greater in region *ii*. Additionally, as discussed above, gene flow will be less. Speciation is thus more strongly favoured in region *ii* than in region *i*, and consequently species richness tends to be higher in region *ii*.

**Figure 7:**
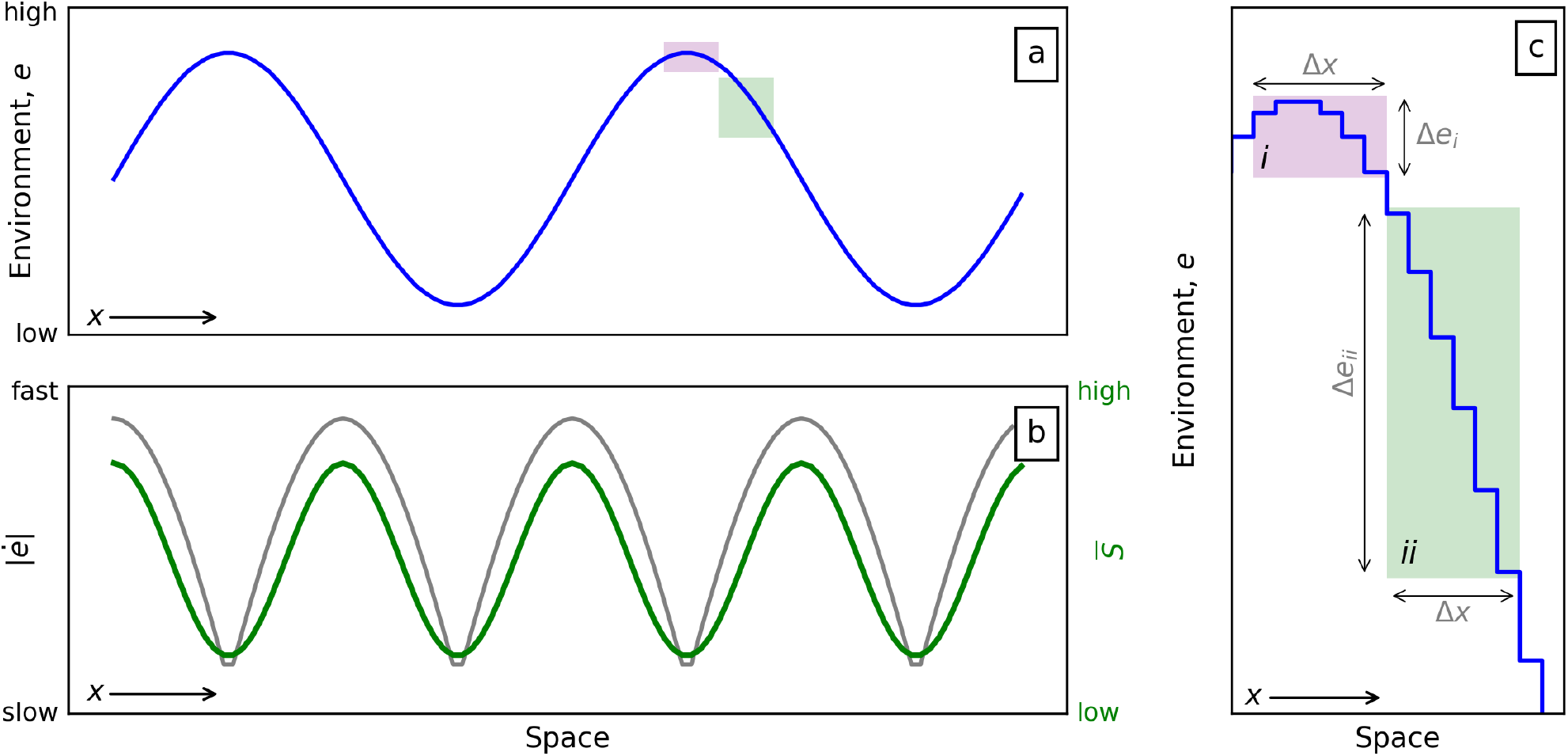
Relationship between environment, its derivative and species richness. (a) Environment, *e*, along the *x*-direction (space; e.g. elevation), see Figure 1. Purple and green boxes refer to the position of *i* and *ii* of panel c. (b) Gray line = absolute derivative of the environment, |*ė*| = |d*e/*d*x*|, which indicates the spatial rate of change of the environment. Green Line = expected species richness. (c) Model parameterisation. Detail from panel a, which shows the discrete values of the simulated environment. Note that Δ*x* is the same for *i* and *ii* and that differences in environment Δ*e*, are larger for *ii* (i.e. Δ*e*_*ii*_ = 61) than *i* (i.e. Δ*e*_*i*_ = 12).

We also find that the coefficient of correlation between |*ė*| and mean species richness 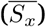 is related to the dispersion ability of organisms (Figure 6). This phenomenon can also be interpreted in terms of gene flow between cells, which is expected to positively correlate with the ability to disperse (Garant et al., 2007). Where dispersion ability is low, gene flow between adjacent cells becomes low, as the probability of an offspring attempting to settle in a different cell to its parents is low (e.g. *P* (*D <* 1 px) = 94% when *d* = 50, see Table 1). As discussed in the previous paragraph, the likelihood of speciation between any pair of cells is controlled by the relative strengths of gene-flow and different adaptive pressures between the cells. When gene flow is low (but greater than zero; e.g. *d* = 35; Figure S3j), speciation between cells will become the favoured outcome in cell-pairs even for those with low adaptive differentials (i.e. in regions with low |*ė*|). When dispersal ability is low therefore, the importance of the magnitude of adaptive differentials between cells becomes less, as most differentials will be sufficiently large to ensure speciation. The correlation between between |*ė*| and 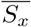 is hence weaker when dispersal ability is low, as variation in |*ė*| is more weakly tied to the prevalence of speciation.

**Table 1:**
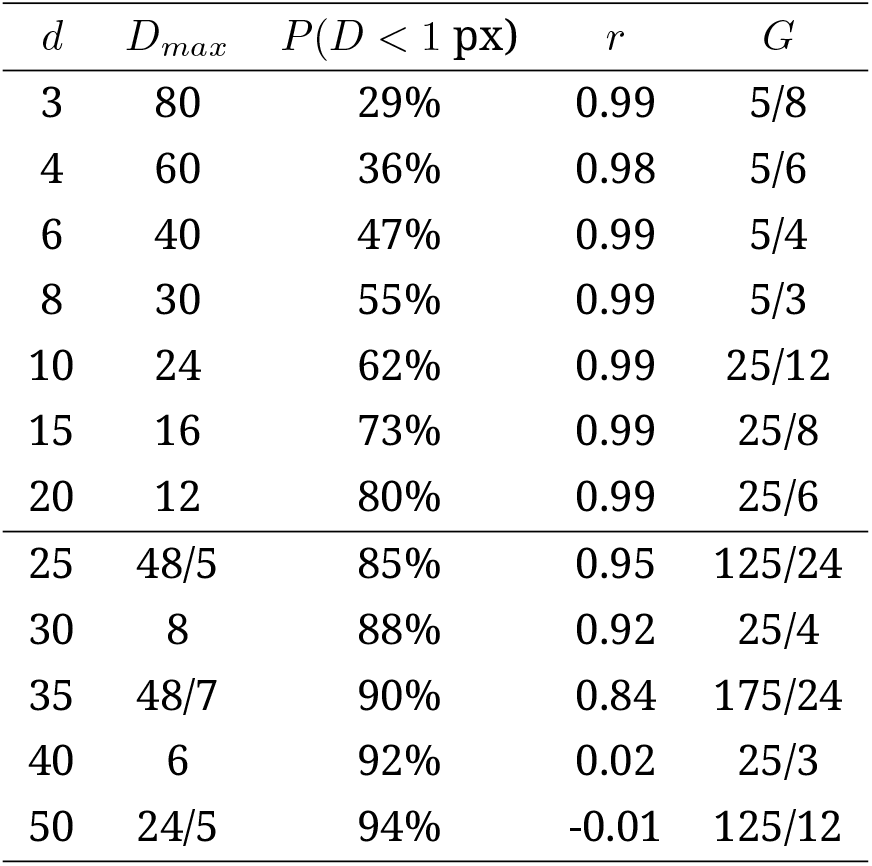
Relationships between dispersion and environmental variability. *d* = dispersion parameter. *D*_*max*_ = 240*/d* = maximum dispersion. *r* = Pearson’s correlation coefficient. *G* = dimensionless dispersion-environment number. Note *λ* = 50.

A feature of our results is that correlation coefficients between |*ė*| and 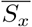 are consistently very high (*r* ≥ 0.99) where *d* is over a particular value (25). The equivalent of this threshold value of *d* cannot be determined for real-world datasets, as our experiments use arbitrary units. To allow real-world comparisons, we define a dimensionless parameter, *G*, that is the ratio of spatial scales to dispersal scales

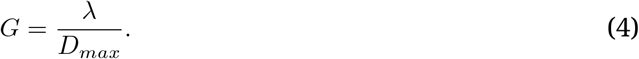

*D*_*max*_ is preferred here over mean displacement as it is, perhaps, more easily measurable in the real-world, and also because it captures the spatial range at which gene flow is possible in each time step. Values of *G* for different dispersion parameters are shown in Table 1. When *G* ≳ 5, |*ė*| is highly correlated with 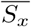.

Where *d* ≥ 40 the correlation coefficients between |*ė*| and 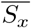 are close to zero (Table 1; Figure 6). We interpret this as an artefact of the integer parameterisation of *e* in our study. As discussed above, in any pair of adjacent cells with differing *e*, speciation will be favoured if differential adaptive pressure arising from the difference in *e* outweighs the homogenisation provided by gene flow. When dispersal ability is low, gene-flow is low, and hence difference in *e* between cells required to favour speciation is also lowered. If dispersal ability is lowered sufficiently, the difference in *e* between cells required to favour speciation may fall beneath 1. Because *e* is an integer, the lowest possible difference in *e* between adjacent cells in the *x*-direction is 1, except where it is precisely zero; in our environment maps, adjacent cells in the *x*-direction with differences of *e* of 0 occur only at minima and maxima. For this reason, in our experiments there must exist a threshold level of dispersal ability below which all cell-pairs in the *x*-direction will favour speciation (except the very few where the difference in *e* is zero), and hence below which the correlation between |*ė*| and 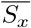 will be very close to zero. We infer that this threshold is between *d* = 35 and *d* = 40. In a real-world dataset, *e* would not be constrained to be an integer, and thus there would be no equivalent threshold level of dispersal ability below which speciation is always favoured. Note that while we infer that the near zero correlations at high *d* are artefactual, we contend that the less precipitous decreases in correlation where *d* ≥ 25 (*G* ≥ 5) are not artefacts, as our explanation for their origin does not depend on integer discretisation. We predict that high correlations will be found in real-world datasets between |*ė*| and 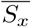 where *G* ≥ 5, and that correlations will become weaker for *G* ≤ 5.

### 4.2 Future work

The digital organisms in our experiments have reproductive and ecological attributes designed to mimic sexual-reproducing sessile organisms such as plants, brachiopods, or corals. Thus, our approach and the results are probably best suited to understanding the species richness of such organisms. The REvoSim model assumes that dispersal capacity is the same for all organisms in each experiment, while this is a simplification, dispersal capacity is frequently observed to be similar within broad clades (Kinlan and Gaines, 2003; Alzate and Onstein, 2022). We note that the frequency distribution of dispersal used in our replicates is similar to some real-world organisms (e.g. trees, Nathan and Muller-Landau, 2000), in that offspring tend to settle closer to parents, with decreasing frequency at increasing ranges. In future work we plan to combine experiments of the type presented here with knowledge of dispersion and environmental variability to establish the diversity distributions of such sessile organisms to establish their variation across scales.

Dispersal ability in our study is constant for each replicate, and identical for all organisms in that replicate. A more nuanced understanding of the relationship between spatial structure and biodiversity may be obtainable by allowing dispersal ability to vary under genetic control. Additionally, environments are invariant over time; permitting them to vary as a function of time as well as space may allow further insights. Both of these extensions are beyond the scope of this baseline study.

## 5 Conclusions

By employing a first-principles model of evolution, which incorporates stochastic dispersion, mutation and a static environment, we established that species richness correlates with the spatial rate of change of the environment. The strength of the correlation is principally dependent on two factors: the dispersion ability of the organisms and the wavelength of the environment. The role of these factors can be understood through established ecological and evolutionary mechanisms. The ratio of environmental wavelength to the maximum dispersion of offspring is proposed as predictor of this correlation in real-world datasets. We suggest that it could be fruitful to investigate relationships between the (spatial) rate of change of environmental variables and species richness across scales in the real world.

## Acknowledgements

Thanks to Francois van Schalkwyk for the HPC support, and Sameer Bansal for the coding review.

