## Supporting Information for "On the relationship between spatial environmental variability, dispersion and biodiversity"

Giulia Bernardini[1a], Gareth G. Roberts[1b], Mark Sutton[1c]

August 15, 2024

[1]Department of Earth Science & Engineering, Imperial College London, Royal School of Mines, Prince Consort Road, London SW7 2BP, UK

[a]**
[b]**
[c]**

This Supplementary Information document contains five figures. Figure S1 is a schematic showing the setup of a simulation, replicates, and the calculation of associated statistical properties.

The remaining figures show how species richness varies as a function of environmental conditions and dispersal ability. They are produced using all replicates for each value of $d$, and show results at the final time step, $T = 100,000$. Figure S2 additionally shows the relationship between species richness and genetic diversity. Note their positive relationship when $d \lesssim 40$. These results are discussed further in the body text of the main manuscript. Figure S3 shows transects through mean species richness and co-located derivatives of the environment. This figure is analogous to Figure 5, but without normalisation. It visually reinforces the view that mean species richness is highly correlated with the derivative of the environment when dispersal ability is high, i.e. $d \lesssim 35$. Figure S4 shows histograms of species richness. We note that when $d \lesssim 25$ that these distributions are skewed towards lower values, and that they are skewed to higher values when $d \gtrsim 25$. Figure S5 shows maps of mean species richness. This figures expands the visualisation of results shown in Figure 4c of the main manuscript. Note that when dispersal ability is relatively high ($d \lesssim 40$) species richness closely follows the derivative of the environment. When dispersal ability is low ($d \gtrsim 35$) similar patterns do not arise. These results are discussed in the main manuscript.

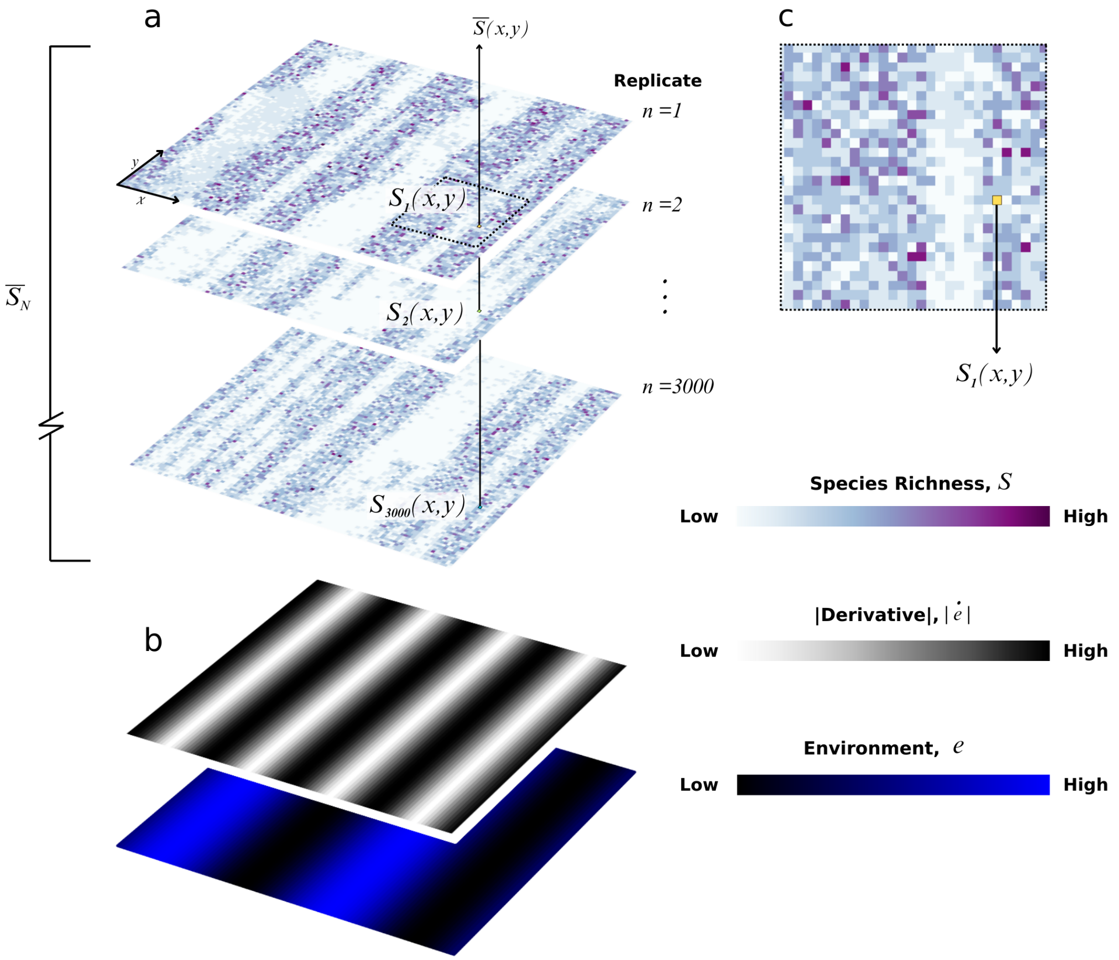


Figure S1: **Schematic showing model setup and associated statistics.** (a) Species richness grids for example replicates for dispersion parameter, $d = 15$. Note $\overline{S_N}$ annotation, i.e. mean species richness for all cells in all replicates. (b) Environment, $e$, grid and its derivative, $|\dot{e}|$. (c) Zoomed area of replicate $n = 1$. Note individual pixel annotated $S_1(x, y)$ (see body text for extended discussion).

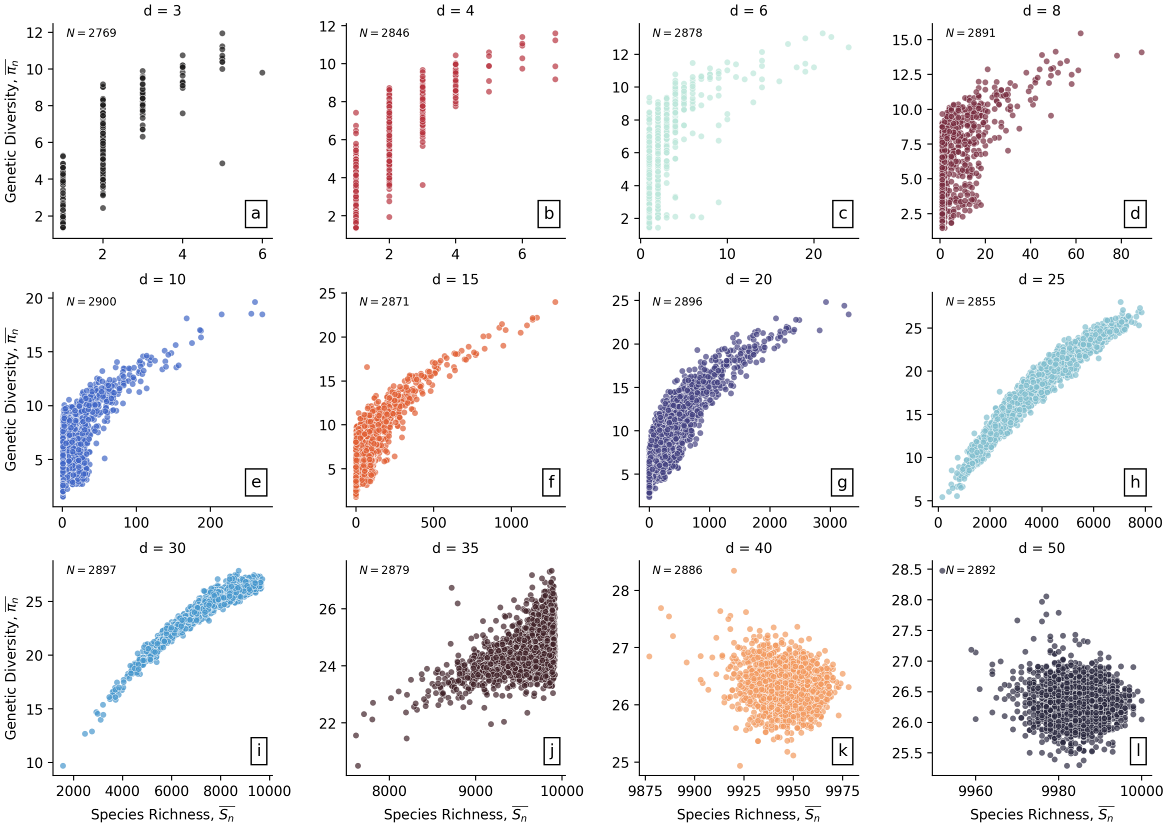


Figure S2: **Genetic diversity versus species richness at equilibrium**. (a) Species richness and genetic diversity compared at $T = 100,000$ for $N$ replicates with dispersion parameter $d = 3$ (see Figure 5a). (b–l) Results for different values of dispersion parameter (annotated).

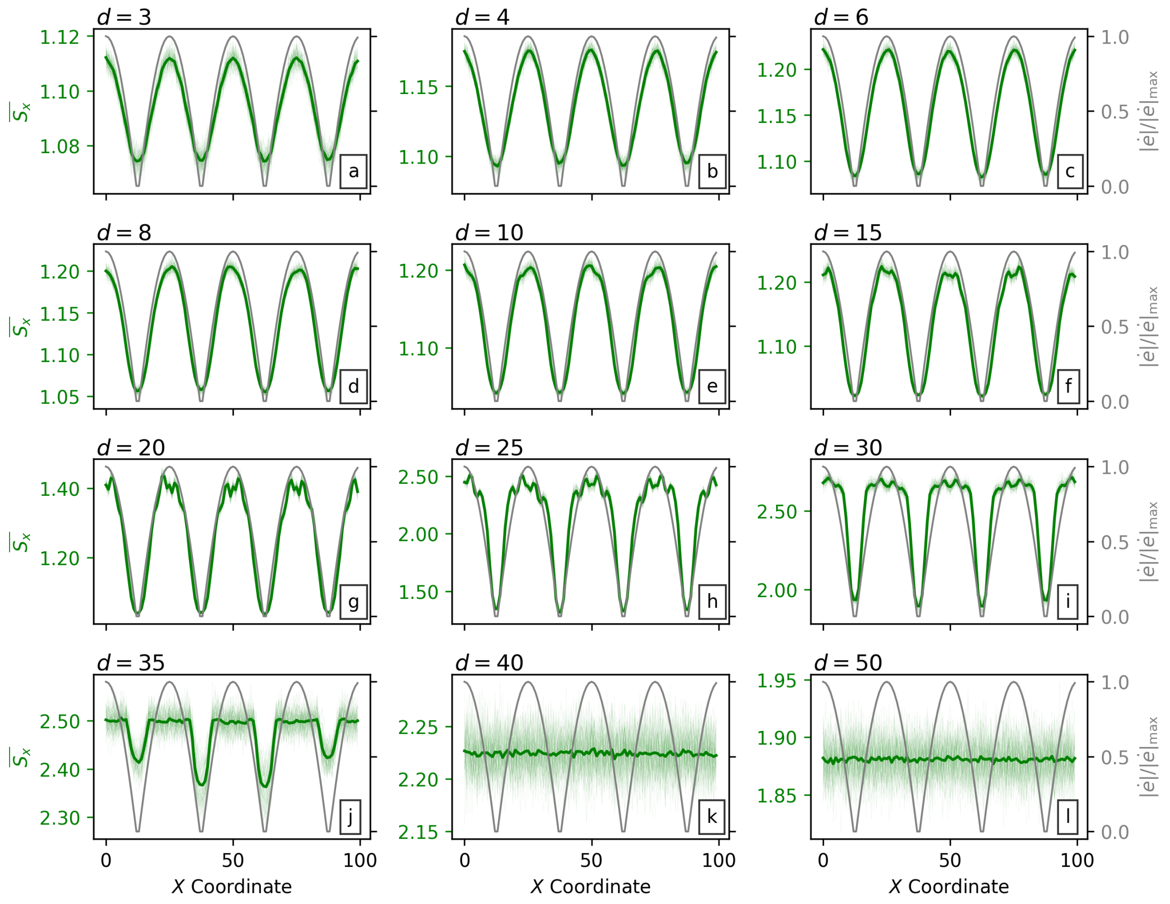


Figure S3: **Relationships between environment and species richness as a function of dispersion and position in $x$-direction**. Same as Figure 5 in the main manuscript, without normalisation of species richness. Gray curves = absolute value of the derivative of the environmental variable, $|\dot{e}|$, normalised by the maximum value, $|\dot{e}|_{\max}$, as a function of $x$ coordinate (see Figures 2a & 4a). Light green curves = mean species richness, $\overline{S}(x, y)$, along transect at $y = 1, 2, \ldots, 100$ from the 3000 simulations. Dark green curves = mean species richness of all 100 transects, $\overline{S}(x)$ (see Figure 4f).

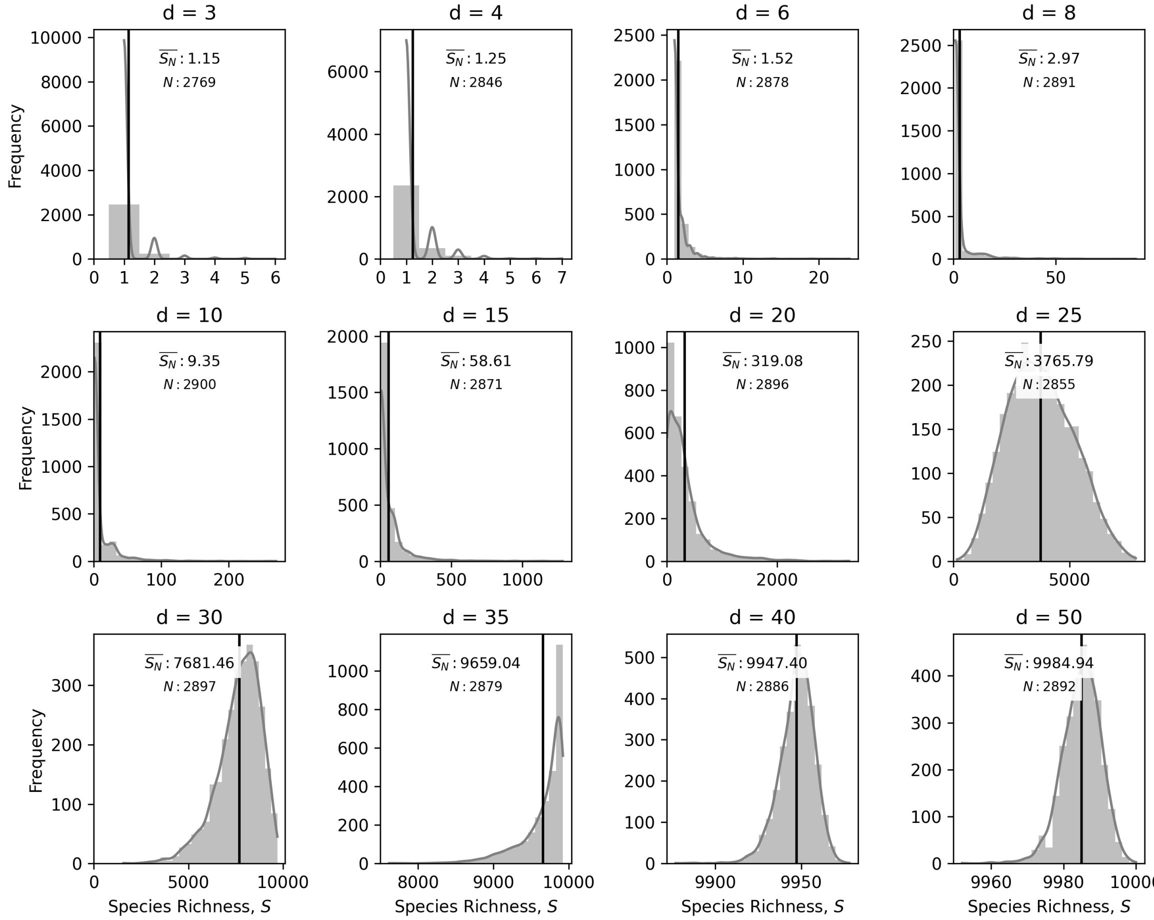


Figure S4: **Distribution of species richness at $T = 100,000$ as a function of dispersion parameter $d$.** Gray bars = frequency of species richness. Black line = mean species richness. Thin dark grey curve = kernel density estimate for species richness. Species richness for $d = 40$ and $d = 50$ is skewed towards $10,000$, which correspond to having unique species in almost all cells in the simulated domain.

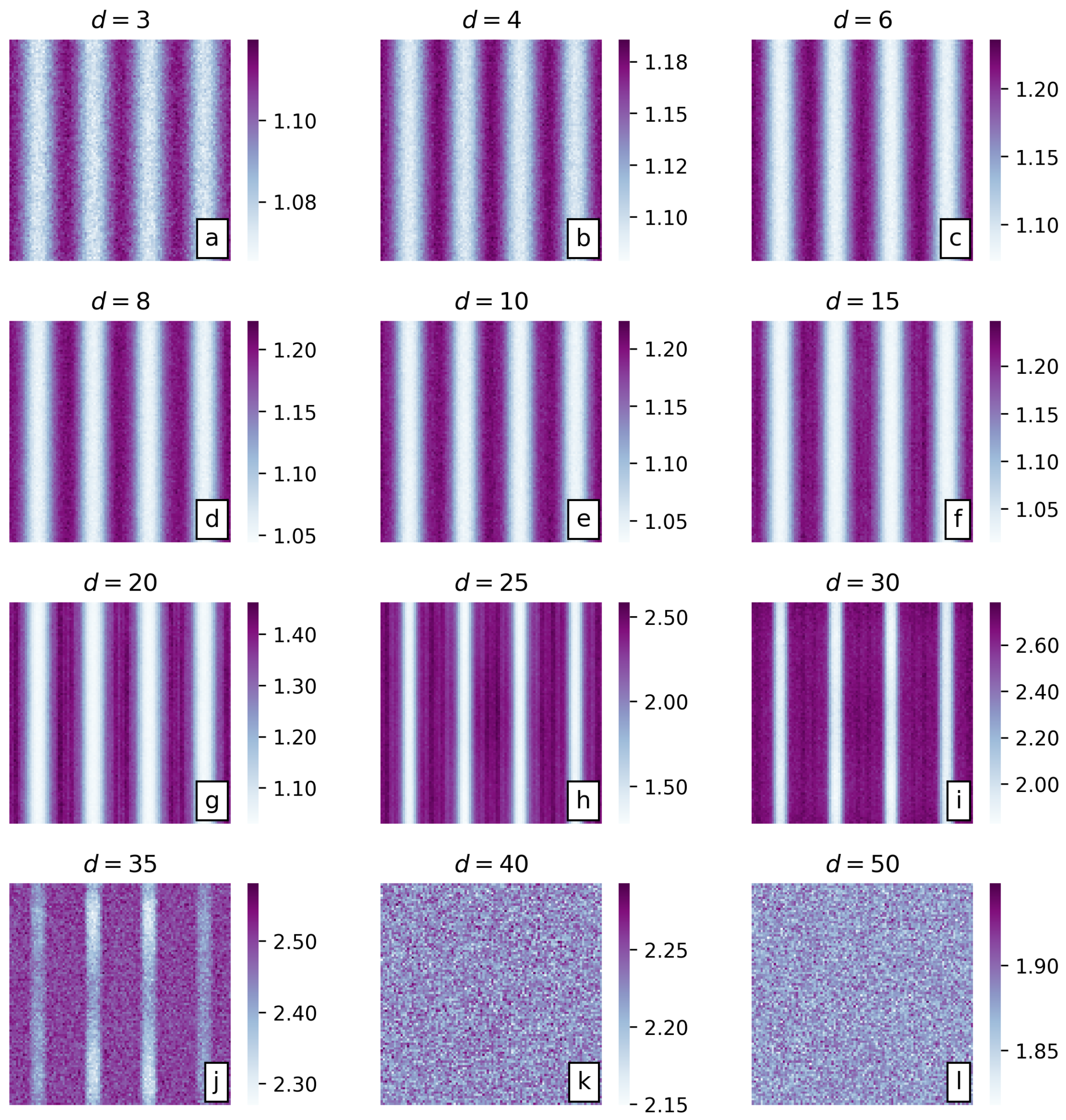


Figure S5: **Mean species richness in each cell at** $T = 100,000$ **from 3,000 simulations for each value of dispersion parameter,** $d$**, tested.** Note that lower values of $d$ indicate greater dispersal ability. The $d$ value with the highest mean species richness per cell, $\bar{S}(x, y) = 2.78$ is $d = 30$ ($D_{max} = 8$ px, $\mu = 0.5$ px). Cyclicity of $\overline{S_x}$ decreases for $d \geq 35$. See body of text for discussion.
